# Intrinsic ignition-based propagation networks reveal hierarchical propagation pathways of spontaneous activity in the human brain

**DOI:** 10.64898/2026.07.16.738814

**Authors:** Jiangling Liu, Xi Chen, Xuhong Liao

## Abstract

Directed communication across brain regions is fundamental to human brain function, yet how local spontaneous activity links to directed whole-brain propagation patterns remains elusive. Prior studies have characterized local propagation events and global wave-like dynamics, but a framework linking local events, interregional propagation pathways, and canonical large-scale propagation patterns remains lacking. Using resting-state functional MRI data from the Human Connectome Project 7T cohort and an intrinsic ignition framework, we quantified cascades of suprathreshold activity events and constructed directed propagation probability networks. We found spatially heterogeneous regional propagation preferences, with high outgoing propagation preferences predominantly localized in somatomotor, visual, and default-mode regions. At the functional system level, propagation modules broadly separated visual, somatomotor, dorsal attention, and ventral attention networks from association systems. Regional stepwise propagation pathways were aligned with the principal functional gradient, and greater hierarchical separation between regions was associated with longer propagation distance. These pathways formed two canonical propagation patterns linking primary and association systems, with somatomotor regions preferentially initiated in bottom-up propagation, default-mode regions in top-down propagation, and attention networks occupying intermediate positions. Notably, the two patterns were not simple mirror images, with top-down propagation showing substantial deviations within somatomotor and visual systems. These findings were replicated in an independent cohort. Moreover, individual propagation architecture predicted cognitive performance and tobacco-use behavior. Collectively, our findings provide cross-scale insights into spontaneous activity propagation in the human brain by bridging local ignition events with hierarchical whole-brain propagation pathways and behaviorally relevant individual differences.

**Significance:** Spontaneous brain activity is often quantified by the temporal synchronization between regions, but this approach does not show how local activity propagates across the brain. We used an interpretable ignition framework to link local spontaneous events and their cascades to directed whole-brain propagation patterns. This approach revealed outgoing propagation preferences of primary and default-mode regions, a modular architecture separating primary and attentional systems from association systems, and two canonical propagation patterns aligned with the brain’s functional hierarchy. These two patterns were not simple mirror images, indicating partially distinct bottom-up and top-down routes. Individual propagation architecture also predicted cognitive performance and tobacco-use behavior. These findings provide a path-based view of spontaneous brain dynamics.

## Introduction

Brain function depends on coordinated interactions and efficient communications among distributed brain regions (1–4). Neuroimaging-based functional connectomics has revealed key principles of human brain network organization by quantifying interregional synchrony in spontaneous activity (5, 6). The functional connectome exhibits non-trivial topological properties, including small-world properties (7–9), modular architecture (10–12), and highly connected hubs (13–15). More recently, gradient-based analyses have revealed a principal axis of functional connectivity organization, with primary sensorimotor systems and default-mode systems located at opposite ends (16–18). This axis has been linked to hierarchical information processing, ranging from sensory perception to multimodal integration and abstract cognition (16–18). It has also provided valuable insights into individual cognition and behavior (16), as well as alterations associated with brain development (19–21) and neuropsychiatric disorders (22, 23). However, this synchrony-based gradient primarily captures a static spatial layout rather than the temporal propagation of activity across regions. How spontaneous activity propagates across the brain and how the dynamics relates to the macroscale functional hierarchy remain to be elucidated.

Accumulating resting-state functional magnetic resonance imaging (R-fMRI) studies indicate that spontaneous brain activity exhibits structured propagation patterns, including systematic time-lag relationships across regions (24, 25), event-triggered cascades (26, 27), and wave-like spreading (28–31). Such propagation dynamics varies with perceptual and cognitive demands (28, 32), developmental stages (33, 34), and brain disorders (35, 36), suggesting that directed activity propagation may play an important role in supporting healthy brain function. Importantly, recent studies have further shown that large-scale activity propagation can be summarized by a small number of dominant spatiotemporal components, such as sensory to association propagation, task-positive to default-mode propagation, and somatomotor to visual propagation (29, 30, 34, 37). However, the local and global descriptions of spontaneous propagation remain insufficiently integrated. Local analyses typically focus on event-level cascades, pairwise delays, or local trajectories (25, 28), whereas global analyses often characterize brain-wide propagation morphology or low-dimensional spatiotemporal modes (29, 30). It remains unknown how local propagation events are routed through specific interregional pathways to give rise to canonical brain-wide propagation patterns.

To address this gap, we employed an interpretable intrinsic ignition framework (38, 39) to construct directed propagation probability networks from spontaneous high-amplitude activity events. Here, we used R-fMRI data from 157 healthy adults from the Human Connectome Project 7T (HCP-7T) (40, 41), and the high spatial resolution of the 7T data is particularly suited to resolving local activity events. By transforming local ignition-event cascades into directed region-to-region propagation probability networks, this approach allowed us to quantify fine-grained interregional propagation pathways and their organization into system-level and whole-brain patterns. We characterized regional propagation preferences, system-level propagation architecture, stepwise propagation pathways, and canonical whole-brain propagation patterns. We further examined how these pathways related to the functional hierarchy and whether individual propagation networks predicted cognitive and behavioral performance. Together, these analyses linked local events, directed interregional pathways, hierarchical cortical organization, and behaviorally relevant individual differences, providing cross-scale insights into spontaneous activity propagation.

## Results

We characterized brain-wide activity propagation based on the intrinsic ignition framework (38, 39), which assumes that high-amplitude ignition events propagate to other regions within a limited temporal window, giving rise to spatiotemporal cascades (Fig. 1A). The cerebral cortex was parcellated into 1,000 cortical regions of interest using the Schaefer 1000-parcel functional atlas (42). After excluding two parcels without valid BOLD time courses, 998 valid cortical nodes were retained for subsequent analyses. Regional ignition events were defined as upward threshold-crossing events in the corresponding regional blood oxygenation-level-dependent (BOLD) time course (Fig. 1B). The propagation of these ignition events was quantified with an interregional propagation probability matrix, in which each element, *p_ij_*, quantified the likelihood that an ignition event in node *i* was followed by a subsequent event in node *j* (Fig. 1C). See *Supplementary Methods* for details.

**Figure 1.**
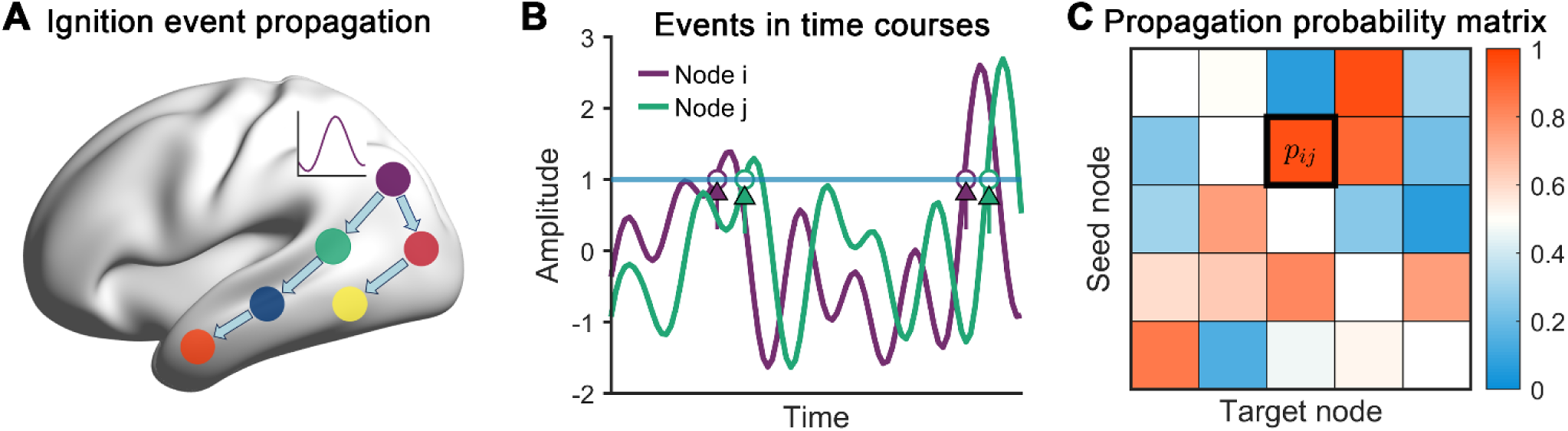
Construction of the propagation probability matrix. (A) Overview of ignition event propagation within the intrinsic ignition framework. (B) Definition of intrinsic ignition events. For each region, an ignition event was identified as an upward crossing of the normalized BOLD time course above a predefined threshold (z = 1). (C) Propagation probability matrix. Each element *p_ij_* quantifies the probability that an ignition event in node *i* was followed by an ignition event in node *j* within a prespecified temporal window (i.e., 5 seconds).

### Regional and functional system-level organization of propagation probability networks

We constructed the group-level propagation probability matrix (Fig. 2A). This matrix exhibited interregional asymmetry (asymmetry index = 0.13; Fig. 2B), indicating modest but measurable directional biases in event propagation. The edge-wise propagation asymmetry was significant but weakly correlated with conventional functional connectivity estimated using Pearson’s correlation (*r* = – 0.015, R^2^ = 2.25×10^-4^, *p* < 0.001). We summarized regional propagation preferences as the ratio of mean outgoing to mean incoming propagation probability. Propagation preferences were spatially heterogeneous across the cortex (Fig. 2C), with high values mainly located in sensorimotor and visual regions, superior parietal lobule, and the precuneus. Given prior functional system definition (42), these propagation preferences differed across systems (Welch’s one-way analysis of variance (ANOVA), *F* (6, 368.91) = 51.86, *p* < 10^-4^, ω^2^ = 0.21). Post hoc comparisons showed higher outgoing preferences in the somatomotor, visual, dorsal attention, and default-mode networks and lower preferences in the ventral attention, limbic, and frontoparietal networks (Games-Howell, false discovery rate (FDR)-corrected; Fig. S1).

**Figure 2.**
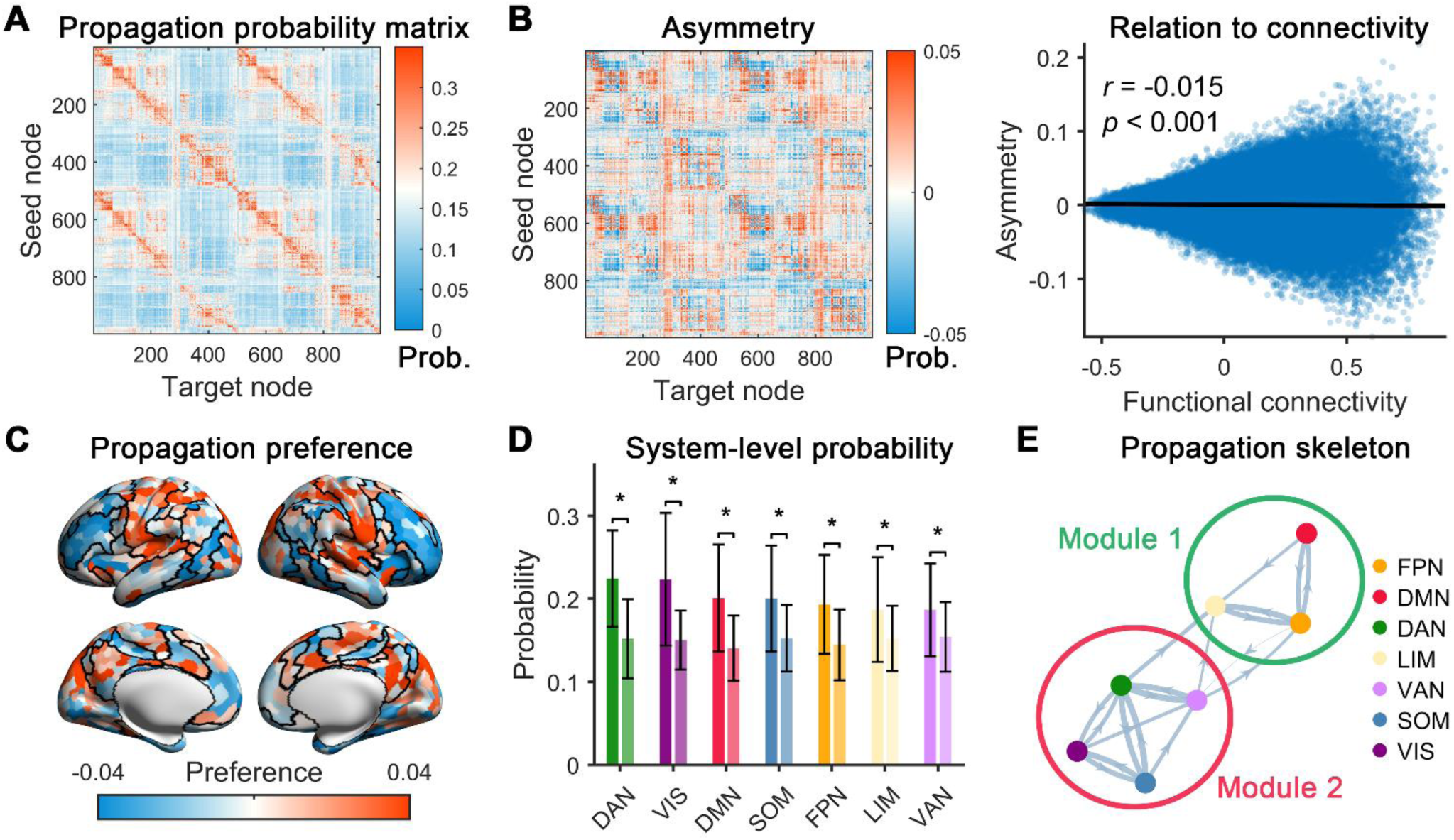
Characteristics of the group-level propagation probability network. (A) Group-level propagation probability network. (B) Asymmetric component of the propagation probability matrix and its relationship to conventional functional connectivity. This component was computed as the elementwise difference between the propagation probability matrix and its transpose. The correlation was calculated using Pearson’s correlation between the corresponding upper-triangular elements of the asymmetric component and the functional connectivity matrix. (C) Spatial distribution of event-count-corrected regional propagation preferences. Propagation preference was defined for each node as the ratio of mean outgoing to mean incoming propagation probabilities and was then residualized by regressing out node-wise intrinsic ignition event counts. (D) Within-system and between-system propagation probabilities. For each system, values are shown as mean ± SD across the corresponding within– and between-system probabilities. (E) Modular organization of the functional system-level propagation skeleton. DAN, dorsal attention network; VIS, visual network; DMN, default-mode network; SOM, somatomotor network; FPN, frontoparietal network; VAN, ventral attention network; LIM, limbic network.

At the functional system-level, propagation probabilities were higher within than between systems (Fig. 2D). The within-system predominance was significant for all seven functional systems (all *p*s < 0.05, Bonferroni-corrected, Fig. 2D). We further constructed a system-level propagation skeleton comprising seven functional systems as nodes by retaining the strongest within– and between-system propagation probabilities while preserving network connectedness (Fig. 2E). This skeleton exhibited a modular structure (*Q* = 0.27), with the visual, somatomotor, dorsal attention, and ventral attention networks forming one module, and the default-mode, frontoparietal, and limbic networks forming the other. Intermodular propagation primarily involved the ventral attention, dorsal attention, frontoparietal, and limbic networks.

### Stepwise propagation reveals fine-grained hierarchical pathways and two canonical patterns

To delineate the fine-grained routes through which ignition events propagate across the cortex, we derived a directed binary propagation network by retaining strong edges in the propagation probability matrix (*p_th_* > 0.2) (Fig. S2; see *Supplementary Methods*). For each region, we calculated the stepwise shortest-path distances, referred to as propagation steps, from this region to all other regions (Fig. 3A). We first assessed whether the overall layout of each region’s stepwise propagation pathway was aligned with the cortical hierarchy captured by the principal functional gradient (see Methods). For each region, we computed Spearman’s correlation between its propagation-step distances to all target regions and the corresponding principal functional gradient scores (Fig. 3B). Regions with positive correlations were primarily located in primary sensorimotor and visual regions as well as the superior parietal lobule, whereas regions with negative correlations were primarily located in the medial and lateral frontal cortices, lateral temporal cortex, angular gyrus, and posterior cingulate cortex/precuneus (Fig. 3B). These correlations were bimodally distributed (Fig. 3C, Hartigan’s dip test, dip = 0.0738, *p* < 0.001), with strong separation between the two modes (Ashman’s *D* = 4.80, see *Supplementary Methods*). At the functional system-level, positive correlations in the somatomotor, dorsal attention, and visual networks indicated stepwise propagation profiles that follow the sensory-to-association axis, whereas negative correlations in the default-mode, frontoparietal, and limbic networks indicated propagation profiles oriented in the opposite direction. Notably, the ventral attention network exhibited near-zero median correlations, suggesting heterogeneous or less consistently gradient-aligned propagation profiles.

**Figure 3.**
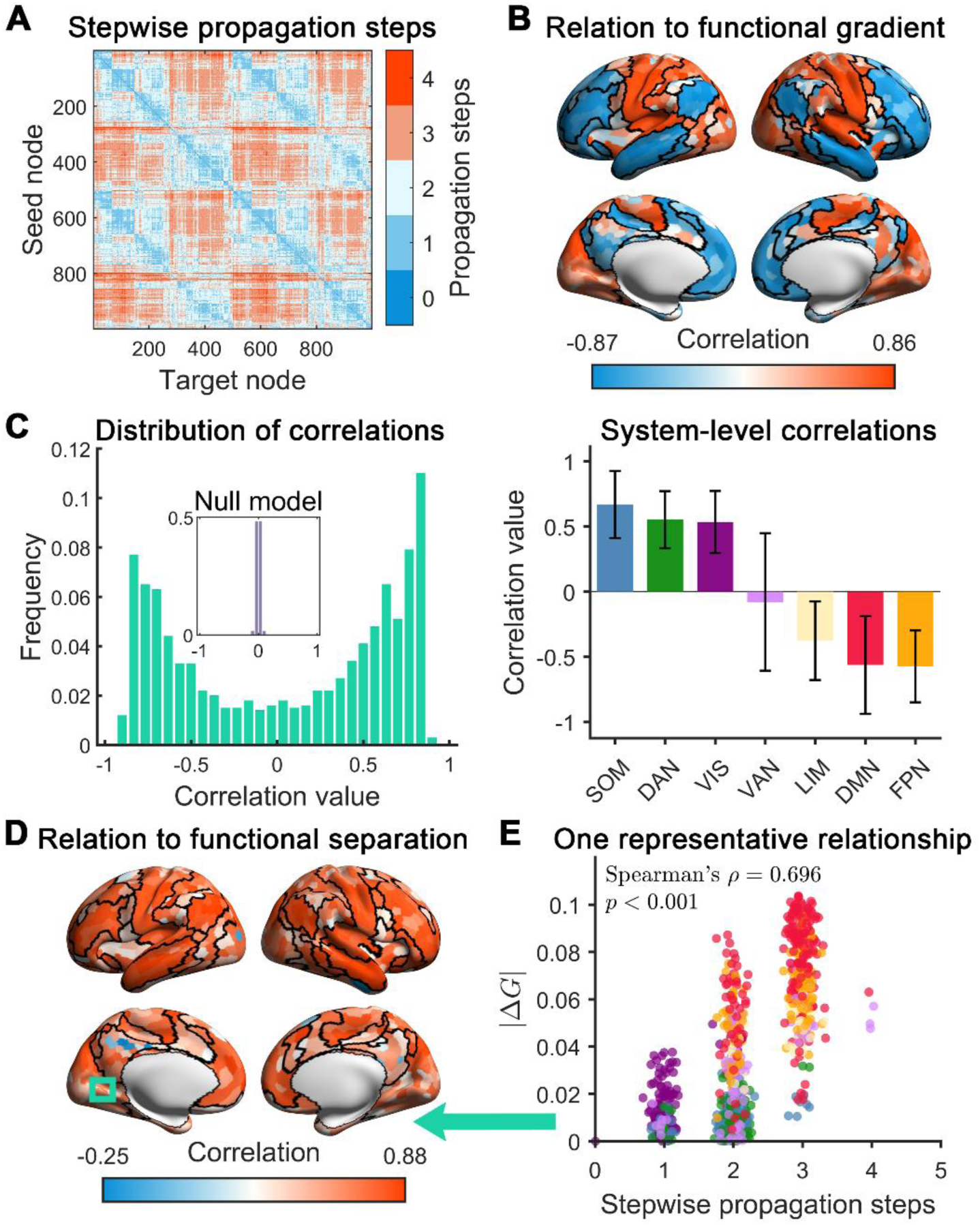
Stepwise propagation distances in relation to the principal functional gradient. (A) Interregional propagation distance matrix. Propagation distances were derived from a binarized group-level propagation probability network obtained by thresholding the matrix with a probability of 0.2 (Fig. S2; see *Supplementary Methods*). (B) Regional alignment between each region’s shortest-path propagation profile and the principal functional gradient. For each seed region, spatial alignment was quantified as Spearman’s correlation between its shortest-path distances to all other regions and the functional gradient scores of the corresponding target regions. (C) Bimodal distribution of seed-wise propagation-gradient correlations and their system-level distribution. The inset shows a null distribution obtained from the aggregation of region-specific permutations of shortest-path distance vectors across target regions (1,000 permutations for each region) (see *Supplementary Methods*). For the system-level distribution, bars indicate the mean correlation across regions within each system, and error bars denote ± 1 SD. (D) Seed-wise associations between propagation distances and hierarchical separation. For each seed region, this association was quantified as Spearman’s correlation between its propagation distances to all other regions and the corresponding absolute differences in principal functional gradient scores. (E) Relationship between propagation distance and gradient difference from a representative region to all other regions. Colors of scatters denote the system affiliations of different target regions. |ΔG| denotes the pairwise absolute difference in principal functional gradient scores. DAN, dorsal attention network; VIS, visual network; DMN, default-mode network; SOM, somatomotor network; FPN, frontoparietal network; VAN, ventral attention network; LIM, limbic network.

We next examined the pairwise relationship between propagation distance and functional hierarchical separation. For most regions (97.37%), propagation-step distances to other regions were positively correlated with the corresponding hierarchical separation, quantified as the absolute differences in the principal functional gradient scores (Fig. 3D). This finding suggests that region pairs farther apart along the cortical hierarchy tend to require more propagation steps. A representative example is shown in Fig. 3E.

Building on these region-wise and pairwise hierarchical relationships, we next sought to identify canonical propagation patterns underlying whole-brain ignition-event propagation by clustering brain regions based on their stepwise propagation profiles. A two-cluster solution was most frequently identified across three clustering validity indices and three clustering algorithms (Fig. S3, see *Supplementary Methods*). Cluster 2 primarily comprised default-mode, frontoparietal, and limbic regions, while cluster 1 comprised regions from other systems (Fig. 4A). Two distinct canonical propagation patterns were identified based on the spatial layouts of the two cluster centroids (Figs. 4B and 4C, left panel). For cluster 1, the canonical propagation pathway was strongly and positively correlated with the principal functional gradient (*r* = 0.924, *p*_spin_ < 0.001; spin-based spatial permutation tests, see *Supplementary Methods*) (Fig. 4B, middle panel). Regions in the dorsal attention, somatomotor, visual, and ventral attention networks were preferentially reached at shorter propagation distances, whereas regions in the limbic, frontoparietal, and default-mode networks were reached at longer propagation distances (Fig. 4B, right panel). In contrast, for cluster 2, the canonical propagation pathway showed a significant but relatively weaker negative correlation with the principal functional gradient (*r* = – 0.885, *p*_spin_ < 0.001) (Fig. 4C, middle panel). Regions in the limbic, frontoparietal, and default-mode networks were preferentially reached at shorter propagation distances, whereas regions in other systems were reached at longer propagation distances (Fig. 4C, right panel). Notably, substantial deviations from the principal gradient were observed within the visual and somatomotor systems (Fig. 4C, middle panel), suggesting partially distinct routes for top-down propagation. Together, these results revealed two canonical propagation patterns broadly consistent with bottom-up and top-down activity propagation along the cortical hierarchy.

**Figure 4.**
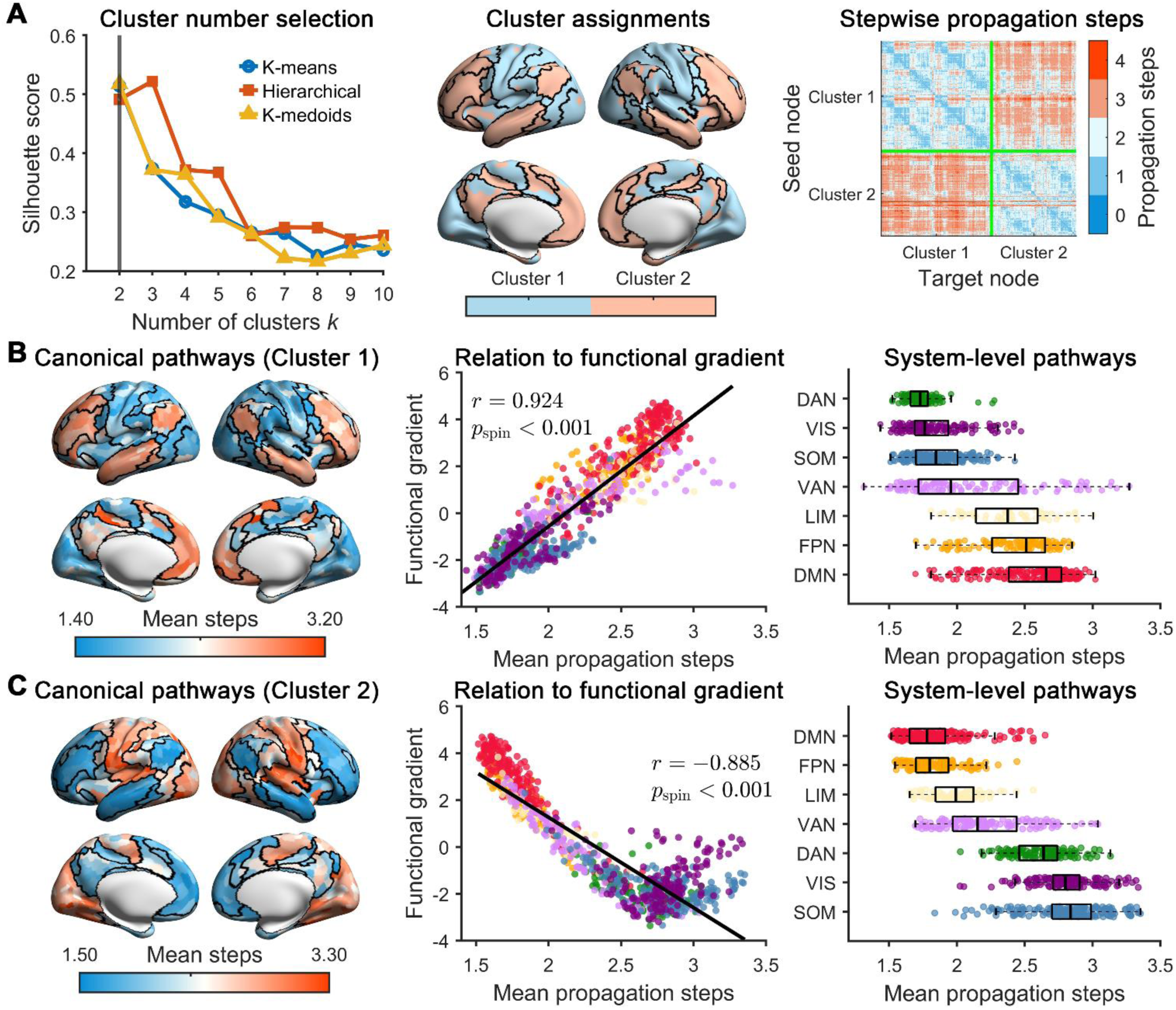
Spatial topography of two canonical shortest-path propagation patterns. (A) Spatial distribution of the two clusters with distinct propagation profiles and the corresponding interregional propagation distance matrix reordered by cluster assignments. The optimal number of clusters (*k* = 2) was most frequently selected across three validity metrics (Silhouette, Calinski-Harabasz, and Davies-Bouldin indices) and three clustering methods (K-means, hierarchical clustering, and K-medoids). (B) Canonical propagation pattern for cluster 1 at the regional and system levels. (C) Canonical propagation pattern for cluster 2 at the regional and system levels. In (B) and (C), canonical propagation distances were derived from the cluster centroids of the regional propagation profiles. Spatial correlations with the principal functional gradient were assessed using spin-based spatial permutation tests (see *Supplementary Methods*). The boxes indicate the interquartile range (IQR) and the median. DAN, dorsal attention network; VIS, visual network; DMN, default-mode network; SOM, somatomotor network; FPN, frontoparietal network; VAN, ventral attention network; LIM, limbic network.

### Individual propagation probability matrices predict cognition and behavior

We next examined whether individual propagation probability matrices could predict cognitive and behavioral performance. Five latent dimensions of cognitive and behavioral scores were considered, including cognition, illicit substance use, tobacco use, personality and emotional traits, and mental health (43). Using support vector regression (SVR) with ten-fold cross-validation, individual propagation probability matrices significantly predicted cognition (*r* = 0.469, *p_perm_* < 0.001; FDR corrected, Fig. 5A) and tobacco-use scores (*r* = 0.387, *p_perm_* < 0.001; FDR corrected, Fig. 5B). For cognitive prediction, high-weight edges primarily involved the prefrontal cortex, precuneus, inferior parietal lobule, lateral frontal cortex, somatomotor areas, and visual regions. At the functional system-level, the high-weight predictive edges were concentrated within the default-mode, somatomotor, and dorsal attention networks, between the somatomotor and dorsal attention networks, and from the default-mode to the visual network (Fig. 5A). For tobacco-use prediction, high-weight edges primarily involved inferior somatomotor regions and the insula. At the functional system-level, predictive contributions were mainly located within the somatomotor network, from the somatomotor network to the ventral attention and frontoparietal networks, and from the dorsal attention to the ventral attention network (Fig. 5B). Notably, the asymmetric distribution of predictive weights across directions indicated that directed propagation probabilities captured behaviorally relevant information, underscoring the value of explicitly modeling propagation directionality. See *Supplementary Methods* for details.

**Figure 5.**
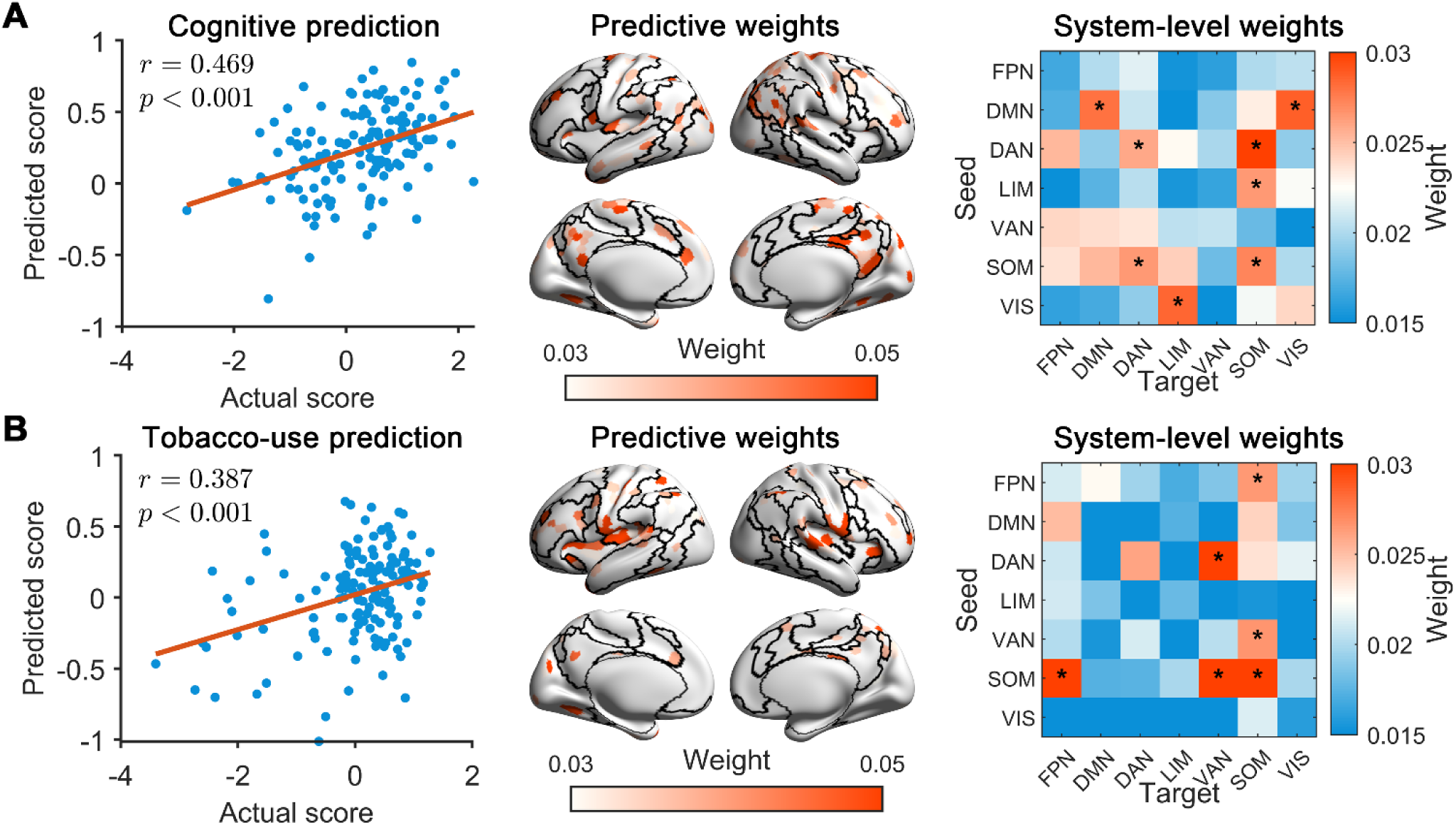
Prediction of cognitive and behavioral performance from individual propagation probability matrices. (A) Prediction of cognitive scores and spatial distributions of prediction weights. (B) Prediction of tobacco-use scores and spatial distributions of prediction weights. In (A) and (B), predictions were performed using support vector regression with ten-fold cross-validation. Statistical significance of prediction performance was assessed with 10,000 permutations, with multiple comparisons corrected using the FDR method. Prediction weights are displayed at the regional and functional system-levels. Asterisks indicate system pairs with weights exceeding the across-pair mean by more than one standard deviation. DAN, dorsal attention network; VIS, visual network; DMN, default-mode network; SOM, somatomotor network; FPN, frontoparietal network; VAN, ventral attention network; LIM, limbic network.

### Robustness and reproducibility of the main findings

We further validated the robustness and reproducibility of the propagation probability structure across different analysis strategies, including i) variations in event and temporal window parameters, ii) hemodynamic delay correction, iii) stringent head motion control, iv) parcellations at different spatial resolutions, and v) validation in an independent Midnight Scanning Club (MSC) dataset (*Supplementary Materials*). Across these validations, the propagation probability matrices, functional system-level propagation skeletons, gradient-related bimodal organization, and two canonical propagation patterns were highly consistent with the main results (Figs. S4-S9). In the independent MSC dataset, despite differences in acquisition parameters and sample composition, we observed similar propagation probability structures and whole-brain propagation patterns (Fig. S9). Collectively, these findings indicate that the observed fine-grained propagation probability structure and canonical propagation patterns are robust to analytical choices and reproducible across independent data.

## Discussion

Using R-fMRI data from two independent datasets, we constructed propagation probability networks and stepwise propagation pathways to characterize the interregional propagation of spontaneous brain activity. Extending beyond previous studies that primarily characterized brain-wide propagation patterns (29, 33, 34, 37), we examined fine-grained propagation profiles at regional and functional system levels. We found that the propagation probability matrix exhibited evident asymmetry and spatially heterogeneous regional preferences. At the functional system level, the propagation skeleton exhibited modular organization, broadly separating primary perceptual and attention systems from association systems. Region-wise stepwise propagation pathways were systematically organized along the principal functional gradient, forming two canonical patterns in which propagation distances increased with hierarchical separation. Finally, individual propagation probability matrices predicted individual cognitive and behavioral performance. Together, these findings reveal structured, directed propagation pathways that link local ignition events to hierarchical whole-brain propagation patterns and behaviorally relevant individual differences.

By leveraging the intrinsic ignition framework (38, 39), we estimated directed, interregional propagation probability matrices. This approach provides a network-level representation of spontaneous activity propagation, linking local high-amplitude events to system-level propagation architecture and gradient-aligned stepwise pathways. In this sense, the present framework complements prior global descriptions of propagating waves, such as time-lag threads, brain-wide wave morphologies, and low-dimensional spatiotemporal modes (25, 29, 30, 33), by specifying which regions preferentially transmit or receive activity and how propagation may proceed through intermediate regions and functional systems.

The propagation probability matrices exhibited robust asymmetry, indicating that spontaneous activity propagation is directionally organized rather than reducible to undirected co-fluctuations. This asymmetry is consistent with the broader notion that communication in brain networks can be directionally biased and spatially heterogeneous (44), although the present analysis focuses on BOLD-dependent propagation rather than direct measurements of neural information transfer. Brain regions exhibited heterogeneous incoming and outgoing preferences even after correcting for local ignition event frequency, suggesting that propagation preferences are not simply attributable to regional differences in ignition event occurrence. Higher outgoing preferences in the somatomotor and visual networks suggest that primary perceptual systems may serve as frequent sources of spontaneous activity cascades, potentially reflecting the broadcast of perceptual or sensorimotor signals (3, 45). In contrast, higher incoming preferences in bilateral frontal, parietal, and temporal cortices suggest that the association regions may act as convergence targets for distributed activity, consistent with their roles in multimodal integration and higher-order processing (15, 45, 46). Interestingly, the precuneus, a well-established structural and functional hub (15), also showed a high outgoing preference. Given its role as a functional core of the default-mode network and its involvement in internally directed cognition and attentional focus tuning (47, 48), this finding suggests that the precuneus may not only serve as a convergent hub for information integration but may also initiate internally oriented propagation cascades.

At the system level, the propagation skeleton revealed that spontaneous propagation is not randomly distributed across functional systems but is organized into a modular architecture. Primary perceptual and attention systems were separated from association systems across analysis strategies and datasets. This separation is broadly aligned with the principal functional gradient, which distinguishes unimodal systems from transmodal association systems (16, 18, 19, 49). This organization may reflect a division between primary perceptual processing and higher-order integrative functions. Importantly, propagation between these modules was bidirectional, supporting both bottom-up propagation from perceptual systems toward association systems and top-down propagation in the opposite direction. The affiliation of dorsal and ventral attention networks with perceptual systems further indicates that attention networks may participate closely in routing sensory-related activity toward higher-order systems. Their involvement near intermodular boundaries may indicate a bridging role in coordinating perceptual processing, attentional selection, and association-level integration, complementing the well-known coordinating role of the salience network in the triple-network model (50).

Building on the regional-level propagation network, stepwise propagation pathway analysis further showed that regional propagation profiles were systematically organized along the principal functional gradient. The bimodal distribution of correlations between regional stepwise propagation profiles and the principal functional gradient suggests that ignition events preferentially propagate along the cortical hierarchy in opposing directions. Previous studies have identified bottom-up and top-down propagation patterns along the cortical hierarchy, using lag-based analysis, optical flow, complex principal component analysis, and related approaches (29, 30, 33, 34). In contrast to these brain-wide descriptions of global propagation patterns, our analysis specifies region-specific stepwise pathways and clarifies how individual regions are embedded within hierarchical propagation routes as a function of hierarchical separation. Primary and default-mode regions, located at opposite ends of the principal gradient, exhibited opposing propagation profiles, consistent with the gradual integration of sensory information and the reciprocal top-down modulation of primary systems (16, 45). Notably, attention systems showed weaker alignment with either direction of propagation. Given their role in linking sensory and transmodal systems (18, 51, 52), their intermediate position may reflect bidirectional information transfer across the cortical hierarchy rather than preferential involvement in either propagation stream.

We identified two canonical patterns of regional propagation profiles. These patterns represent distinct stepwise propagation pathways corresponding to the bimodal propagation tendencies. The bottom-up pattern showed a stronger correspondence with the principal functional gradient, consistent with the unimodal-to-transmodal hierarchy (16–18). In contrast, the top-down pattern showed a relatively weaker gradient association, with prominent deviations in visual and somatomotor systems. This asymmetry suggests that bottom-up and top-down propagation may not simply represent mirrored images along the same route but instead reflect partially distinct propagation mechanisms. Notably, previous functional gradient studies have shown that visual and somatomotor systems, although both located at the unimodal end of the principal gradient, are separated along secondary gradients (16, 18). Our findings suggest that this sensorimotor differentiation may be particularly relevant to top-down modulation, which appears to engage more system-specific feedback pathways. Across both canonical propagation patterns, the ventral attention network occupied an intermediate position between primary systems and association systems, consistent with the system-level modular architecture and supporting a potential role as a bridge for bidirectional propagation. Such a bridging position may allow the ventral attention network to mediate both the relay of perceptual information toward higher-order systems (52, 53) and the redistribution of top-down signals back toward sensory and motor systems (54, 55).

Propagation probability also captured behaviorally relevant individual differences, extending previous work based largely on static functional connectivity. High predictive weights for cognitive performance were widely distributed across multiple functional systems, suggesting that cognition is supported by coordinated, distributed propagation across the brain. This pattern is consistent with connectome-based prediction studies showing that individual cognitive differences are encoded in whole-brain connectivity profiles rather than isolated regional or network effects (56–58). The involvement of the default-mode network suggests that cognitive performance may also depend on internally generated representations, memory-guided simulation, semantic processing, and the integration of internal and perceptual information (59, 60). The contributions of dorsal attention, ventral attention, and limbic systems are consistent with their roles in goal-directed attentional selection, attention orienting to salient events, and emotion-related cognitive modulation (50, 61, 62). The involvement of visual and somatomotor systems indicates that individual cognitive performance may depend on how perceptual and action-related representations are dynamically integrated with attentional and association systems. Notably, for the frontoparietal system, the relatively higher predictive weights for incoming propagation than for outgoing propagation may suggest that frontoparietal contributions to cognitive performance are expressed less as a primary source of spontaneous propagation and more as a convergence or integration stage that receives attentional signals.

Predictive features for tobacco-use performances were primarily related to propagation within the somatomotor network, consistent with previous studies suggesting that somatomotor and motor-related regions are involved in nicotine dependence and habitual smoking behaviors (63–65). These regions have been repeatedly implicated in habit formation and cue-action coupling, reflecting a shift from goal-directed to automatic behavioral control (66). Additionally, we found that high predictive weights were associated with propagation from the somatomotor network to the ventral attention and frontoparietal networks. The ventral attention network and insular regions are known to support salience detection, interoceptive processing, and stimulus-driven reorienting (52, 61), whereas frontoparietal systems contribute to executive control and goal-directed regulation (67, 68). Our findings further indicate that tobacco use may be associated with the routing of sensorimotor-driven signals into salience and executive networks. Together, these findings imply that tobacco-use traits may relate to altered propagation pathways linking habit-related sensorimotor systems with salience and control networks.

Several issues warrant further consideration. First, propagation probability was estimated from BOLD signals and it may be affected by interregional differences in hemodynamic responses (69). Previous electrophysiological and multimodal studies have provided important evidence that large-scale fMRI propagation patterns have a neural basis (25, 30). Our main findings were robust to corrections for regional hemodynamic latencies, further supporting the stability of the observed propagation organization. Future work should validate these pathways using imaging modalities that more directly measure neural activity (3, 70). Second, the limited temporal resolution of fMRI constrains the timescale at which propagation can be resolved. Higher temporal resolution recordings, such as magnetoencephalography (MEG), electroencephalography (EEG), or intracranial electrophysiology, may help characterize propagation at finer timescales (71, 72). Third, dynamic propagation likely depends on the underlying structural architecture (73, 74). It remains unclear how white matter connectivity and local microstructure constrain directional propagation preferences, modular propagation organization, and hierarchical routing. Fourth, we focused on resting-state activity propagation in healthy adults. Extending this framework to task conditions, developmental cohorts, and psychiatric populations may clarify how propagation pathways vary with cognitive demand, maturation, and pathology. Finally, stepwise shortest paths provide an interpretable representation of propagation routes, but they should be viewed as network-derived pathway estimates rather than direct observations of physical signal transmission. Future studies combining perturbation, longitudinal designs, and multimodal recordings will be important for testing the physiological interpretation of these pathways.

## Materials and Methods

### Neuroimaging data and participants

Neuroimaging data were obtained from two resting-state fMRI datasets. The Human Connectome Project 7T dataset (HCP-7T; S1200 release) (40, 41) was used as the discovery cohort and the Midnight Scan Club (MSC) (75) dataset as an independent validation cohort. Both datasets were obtained through established data-sharing repositories. For the HCP-7T dataset, the original study procedures were approved by the Institutional Review Board of Washington University in accordance with the original study protocols. The MSC dataset was collected under the ethical approvals and informed-consent procedures of the original study. Data from 157 healthy young adults (22.0-36.0 years, 66 males) were selected from the HCP-7T dataset. These participants completed four 16-min resting-state runs (64 min in total) and met the HCP quality-assurance criteria. The MSC dataset included densely sampled resting-state fMRI data from ten healthy adults. One participant was excluded because of excessive head motion and sleep during scanning, leaving nine participants for validation analyses. Further details on inclusion criteria and participant demographics (age and sex) are provided in *Supplementary Methods*.

### Construction of propagation probability networks and network topology analyses

Interregional propagation probability networks were constructed from resting-state fMRI data using the intrinsic ignition framework (38, 39) (Fig. 1). From these networks, we quantified regional propagation preference, functional system-level propagation skeletons, and region-specific stepwise propagation distances. A principal functional gradient was derived from group-averaged functional connectivity matrix (16–18) and served as a reference axis for relating propagation organization to cortical hierarchy. Canonical whole-brain propagation patterns were identified by clustering regional stepwise shortest-path profiles, and cluster solutions were evaluated using complementary validity criteria. Further methodological details are provided in *Supplementary Methods*.

### Propagation-based cognitive and behavioral prediction

Brain-behavior associations were assessed using support vector regression with 10-fold cross-validation to predict cognitive and behavioral scores from individual propagation probability networks. Statistical significance was assessed using permutation testing. Prediction weights were summarized at regional and functional system levels (see *Supplementary Methods* for details).

## Data and code availability

The HCP-7T dataset is available through the Human Connectome Project Young Adult S1200 release (https://www.humanconnectome.org/study/hcp-young-adult/data-releases). The Midnight Scan Club dataset is available through OpenNeuro (https://openneuro.org/datasets/ds000224). The Schaefer cortical parcellations, including the 200-, 400-, and 1000-parcel resolutions and their corresponding functional system annotations, are publicly available from the CBIG repository (https://github.com/ThomasYeoLab/CBIG/tree/master/stable_projects/brain_parcellation/Schaefer2018_LocalGlobal).

Publicly available toolboxes used in this study include the Brain Connectivity Toolbox (https://github.com/brainlife/BCT), BrainNet Viewer (https://www.nitrc.org/projects/bnv), and the rsHRF toolbox (https://github.com/compneuro-da/rsHRF). Statistical analyses, clustering, and support vector regression were performed using MATLAB built-in functions. Custom MATLAB scripts and functions developed for propagation probability matrix construction, propagation characteristic analyses, and cognitive and behavioral prediction are available in a Github repository (https://github.com/liaolab-bnu/Propagation_rfMRI).

## Supporting information

SI Materials and Methods for Liu_Manuscript20260716

## Acknowledgements

We thank Saiwen Yu for technical assistance and Dr. Ye Tian for generously sharing cognitive and behavioral scores. We also thank the investigators and participants in the Human Connectome Project and the Midnight Scan Club for their contributions to data collection, curation, and public sharing. This work was supported by the National Natural Science Foundation of China (No. 81971690) and the Tang Scholar Award of Beijing Normal University. The authors declare that DeepSeek-V4 was used solely for language editing. All AI-assisted edits were reviewed and verified by the authors.

