## Supplementary material for "Intrinsic ignition-based propagation networks reveal hierarchical propagation pathways of spontaneous activity in the human brain": SI Materials and Methods for Liu_Manuscript20260716

### *Supplementary Information*

### Contents

**Supplementary Methods**

Datasets

Data preprocessing

Construction of the propagation probability networks

Characterization of intrinsic ignition events and propagation probability networks

Analysis of regional stepwise propagation pathways and their relationship to the cortical hierarchy

Identification of canonical propagation patterns

Prediction analysis of individual cognitive and behavioral performance

Validation analysis

**Supplementary Results**

Validation results

**Supplementary Figures**

**Supplementary References**

### Supplementary Methods
### Datasets

We employed two publicly available resting-state functional MRI (R-fMRI) datasets, the Human Connectome Project 7T (HCP-7T) dataset (1, 2) and the Midnight Scan Club (MSC) dataset (3). The HCP-7T data were used for the main analyses, whereas the MSC data were used for independent validation.

*HCP-7T data*. We used 7T R-fMRI data from the S1200 release of the HCP Young Adult Project (1, 2). MRI data were acquired across four sessions using a 7T Siemens MAGNETOM system with a 32-channel head coil at Washington University in St. Louis. During each session, R-fMRI data were acquired while participants maintained their eyes open and relaxed fixation. Each session included one 16-minute run, yielding four R-fMRI runs (64 min in total) per participant. The R-fMRI data were obtained using a 2D gradient-echo echo-planar imaging (EPI) sequence, with the following parameters: multiband acceleration factor = 5, repetition time (TR) = 1000.0 ms, echo time (TE) = 22.2 ms, flip angle = 45°, matrix size = 130 × 130, field of view = 208 × 208 $mm^2$, 1.6-mm isotropic voxels, and 85 slices. Phase-encoding directions alternated between posterior-to-anterior and anterior-to-posterior across runs. The original study procedures were approved by the Institutional Review Board of Washington University. Following quality control, we excluded participants with excessive head motion in any run (translation or rotation > 3.0 mm or 3˚, or mean framewise displacement (FD) > 0.50 mm). The final sample included 157 healthy adults (22.0-36.0 years; 66 males and 91 females) with complete data across all four runs.

*MSC data***.** The MSC dataset comprised densely sampled R-fMRI data from ten healthy, right-handed adults (24.0-34.0 years; 5 females) (3). MRI data were acquired using a 3T Siemens Trio MRI scanner at Washington University in St. Louis. Each participant completed ten separate late-night sessions, each beginning around midnight. During each session, a 30-minute resting-state run (818 volumes) was acquired, yielding approximately 5 hours of resting-state data per participant. R-fMRI data were collected using a gradient-echo EPI sequence (TR = 2.2 s, TE = 27.0 ms, flip angle = 90°, voxel size = 4 × 4 × 4 $mm^3$, and 36 slices). The dataset was collected under the ethical approvals and informed-consent procedures of the original study. One participant was excluded because of excessive head motion and self-reported sleep during scanning (3, 4), leaving nine participants for validation analyses. The MSC data were used to assess the robustness and generalizability of the group-level propagation features.

### Data preprocessing

*HCP-7T data***.** We leveraged R-fMRI data processed with the HCP minimal preprocessing pipeline adapted for 7T acquisitions (1, 2), including gradient-distortion correction, EPI distortion correction, head motion correction, bias-field correction, intensity normalization, and mapping to CIFTI grayordinates based on multimodal surface matching (MSMAll). The data were further denoised using ICA-FIX, high-pass filtered at 0.009 Hz, and spatially smoothed on the cortical surface with a Gaussian kernel (full width at half maximum = 2.0 mm). In our study, we additionally performed temporal band-pass filtering (0.01-0.08 Hz) to vertex-level time courses.

*MSC data*. We used R-fMRI data preprocessed using the MSC pipeline (3), including slice-timing correction, head-motion correction, bias-field correction, and intensity normalization. Functional data were then registered to each participant's structural images and mapped to CIFTI grayordinates in fs_LR space. Nuisance regression was performed to remove motion parameters and their derivatives, as well as white-matter and cerebrospinal-fluid signals. The resulting data were high-pass filtered at 0.009 Hz. We additionally applied temporal band-pass filtering (0.01-0.08 Hz) to vertex-level time courses, to ensure consistency with the HCP-7T analyses.

**Construction of the propagation probability networks**

Using the intrinsic ignition framework (5, 6), we characterized brain-wide propagation patterns of spontaneous activity by quantifying directed interregional propagation probabilities of intrinsic ignition events (Figs. 1A and 1B). Specifically, the cortical surface was divided into 1,000 nodes based on a prior functionally defined atlas (7), designed to maximize functional homogeneity within each nodal region. After excluding two parcels without valid BOLD time courses, 998 valid cortical nodes were retained for subsequent analyses. For each participant and run, nodal BOLD time courses were extracted by averaging vertex time courses within each node and then normalized into z-score values with zero mean and unit variance (Fig. 1B).

Ignition events were defined as upward crossings of nodal time courses above a predefined threshold (z = 1.0). This procedure yielded a binary event sequence for each node and a corresponding event count $N_i$ for node $i$ within each run. For each ordered node pair $(i, j)$, we computed $C_{ij}$ as the total number of events in node $j$ that occurred within a fixed temporal window following all events in node $i$. When multiple ignition events in node $j$ fell within a given window, all such events were accumulated in $C_{ij}$. The directed propagation probability from node $i$ to node $j$ was defined as $p_{ij} = C_{ij}/N_i$. For each participant, we computed a propagation probability matrix (*PPM*) separately for each R-fMRI run and then averaged the resulting matrices across runs to obtain an individual-level matrix. Group-level propagation probability matrices were subsequently obtained by averaging individual-level matrices across participants.

The temporal window for estimating interregional propagation probabilities was set to 5 s (5 TRs; TR = 1.0 s) for two reasons. First, because the BOLD time courses were band-pass filtered at 0.01-0.08 Hz, the period corresponding to the upper frequency bound is approximately 12.5 s; thus a 5-s window captures approximately 40% of this shortest retained oscillatory timescale. Second, prior time-lag analyses of R-fMRI have reported interregional delays on the order of several seconds (8, 9).

**Characterization of intrinsic ignition events and propagation probability networks**

To visualize directional bias in ignition event propagation, we first characterized the asymmetric component of the propagation probability matrix by calculating the difference between the group-level propagation probability matrix and its transpose. To further quantify the overall degree of asymmetry, we computed an asymmetry index after setting the diagonal elements to zero, defined as:

$$\frac{\|PPM-PPM^T\|_F}{\|PPM\|_F},$$

where $PPM$ denotes the group-level propagation probability matrix and $\|*\|_F$ denotes the Frobenius norm. A value of zero indicates a perfectly symmetric propagation probability matrix, whereas larger values indicate stronger directional bias in interregional propagation. We further compared the spatial similarity between the asymmetric component and the traditional functional connectivity matrix to assess whether the observed directional asymmetry could be explained by interregional synchrony. The functional connectivity matrix was computed using Pearson's correlation, and edge-wise similarity was estimated across the corresponding upper-triangular elements of the two matrices.

Next, we assessed the propagation preference of each nodal region as the ratio of its mean outgoing to mean incoming propagation probabilities. To reduce the potential influence of regional variation in intrinsic ignition event counts, propagation preference was residualized by regressing out node-wise intrinsic ignition event counts. Positive and negative residuals indicated stronger outgoing and incoming propagation preferences, respectively. To test whether these propagation preferences varied across functional systems, nodes were grouped into seven functional systems based on a previously defined functional atlas (10). Welch's one-way analysis of variance (ANOVA) was used to assess the main effect of functional system, given its robustness to unequal variances and sample sizes (11). If the functional system effect was significant ($p < 0.05$), pairwise comparisons were performed using Games-Howell tests (12). Statistical significance of these intersystem comparisons was corrected for 21 (i.e., 7×6/2) comparisons using the false discovery rate (FDR) procedure (13).

We further characterized propagation organization at the functional system-level. First, we compared within- and between-system propagation probabilities in the group-level propagation probability matrix, using Welch's two-sample t-tests. For each functional system, within-system edges were defined as directed edges among nodes within the same system, whereas between-system edges included both outgoing and incoming directed edges linking that system to all other systems. Statistical significance across seven system-wise comparisons were corrected using the Bonferroni correction. Second, we constructed a functional system-level propagation skeleton network, comprising seven nodes representing functional systems. We retained the strongest within- and between-system propagation probabilities while preserving the minimum set of edges required to maintain network connectedness. The spectrum algorithm (14-16) was applied to identify the modular structure in this skeleton network, and the corresponding modularity index ($Q$) was computed (17).

**Analysis of regional stepwise propagation pathways and their relationship to the cortical hierarchy**

Based on the group-level propagation probability network, we identified regional stepwise propagation pathways and corresponding propagation distances for each seed node. Specifically, propagation distances were derived from a binarized group-level propagation probability network by retaining edges with probabilities greater than or equal to 0.20. This threshold preserved network connectedness while retaining as few edges as possible. For each seed node, the shortest-path distances to all other nodes were computed on the binary network using Dijkstra's algorithm (18), yielding seed-specific stepwise propagation profiles. We then assessed whether regional stepwise propagation pathways were systematically organized along the cortical hierarchy. The cortical hierarchy was quantified using the principal functional gradient (19-21), estimated from the group-level functional connectivity matrix. Specifically, for each participant and R-fMRI run, we calculated a functional connectivity matrix using Pearson's correlations between nodal time courses. The group-level functional connectivity matrix was obtained by averaging the resulting matrices across runs and then across participants. Before gradient estimation, diagonal elements and negative correlations in the group-level matrix were set to zero. Principal component analysis was applied to this thresholded matrix, and the first principal component was used as the principal functional gradient (22).

We further assessed the spatial alignment between nodal propagation pathways and the cortical hierarchy. For each seed node, we computed Spearman's correlation between its shortest-path distances to all target nodes and the principal functional gradient scores of the target nodes. This yielded one propagation-gradient correlation coefficient for each seed node. Positive and negative coefficients indicate opposite relationships between stepwise propagation distance and hierarchical position along the principal gradient, thereby characterizing seed-specific propagation orientations along the principal functional gradient.

To account for spatial autocorrelation, the statistical significance of the observed spatial correlation between the canonical propagation patterns and the principal functional gradient was evaluated using spin-based spatial permutation tests (23, 24). Parcel centroids on the fsaverage spherical surface were randomly rotated within each hemisphere and reassigned using one-to-one Hungarian matching (25, 26). Spearman's correlation was recalculated after each of 10,000 rotations to generate a spatial null distribution. To assess the bimodality of these correlation coefficients, we performed Hartigan's dip test for unimodality (27), for which the null hypothesis assumes a unimodal distribution. The dip statistics and its statistical significance were estimated using a Monte Carlo procedure with 5,000 iterations. To further quantify the separation between the two modes, we fitted a two-component Gaussian mixture model to the correlation coefficients using MATLAB's fitgmdist function and computed Ashman's $D$, defined as $D = |\mu_1 - \mu_2| / \sqrt{(\sigma_1^2 + \sigma_2^2)/2}$, where $\mu_1$ and $\mu_2$ denote the component means and $\sigma_1$ and $\sigma_2$ denote the corresponding standard deviations (28). Larger values of Ashman's $D$ indicate stronger separation between the two components.

To assess whether the observed propagation-gradient associations could arise by chance, we generated a null distribution by randomly shuffling the seed-specific propagation profiles across target regions. Specifically, for each seed node, we performed 1,000 permutations, randomly shuffling its shortest-path distances across target nodes in each permutation. We then calculated Spearman's correlation between each shuffled propagation profile and the principal functional gradient, yielding 1,000 surrogate correlations for each seed node. These surrogate correlations were pooled across seed nodes to form a null distribution of propagation-gradient correlations expected when propagation distances are unrelated to the spatial organization of the cortical hierarchy.

To test whether propagation-gradient correlations differed across functional systems, we grouped nodes according to their functional system assignments. The effect of functional system was evaluated using Welch's one-way ANOVA (11). When the system effect was significant ($p < 0.05$), we performed pairwise comparisons using Games-Howell tests (12). The resulting *p*-values were corrected for 21 (i.e., $7 \times 6/2$) comparisons using the FDR procedure.

In addition to examining regional propagation profiles, we further assessed whether interregional propagation distances scaled with hierarchical separation along the principal functional gradient. For each seed node, we calculated Spearman's correlation between its shortest-path distances to all target nodes and the corresponding absolute differences in principal functional gradient scores between the seed node and each target node. Positive correlations indicated longer shortest path distances with increasing hierarchical separation.

**Identification of canonical propagation patterns**

To identify shared structures underlying the regional stepwise propagation pathways, we clustered seed regions using their outgoing shortest-path propagation profiles across target regions as features. Clustering was first performed using k-means and then repeated using hierarchical clustering and k-medoids clustering to assess robustness. The optimal number of clusters was selected by jointly considering three validity indices: the Silhouette score (29), the Calinski-Harabasz index (30), and the Davies-Bouldin index (31). Across the nine method-index combinations, a two-cluster solution was most frequently identified, occurring in seven combinations. For the two-cluster solution, we further quantified the consistency of nodal assignments across three clustering methods using the adjusted Rand index (32).

For each cluster, we obtained the canonical shortest-path propagation pattern as the spatial profile represented by the cluster centroid. We further assessed the spatial alignment of each canonical propagation pattern with the principal functional gradient. To characterize the system-level organization of each canonical propagation pattern, the seven functional systems were ordered by their mean propagation distances, from shortest to longest.

**Prediction analysis of individual cognitive and behavioral performance**

We further assessed whether individual propagation probability matrices contained information related to individual differences in cognitive and behavioral performance. We used linear support vector regression (SVR) with 10-fold cross-validation to predict five cognitive and behavioral dimensions (33), including cognition, illicit substance use, tobacco use, personality and emotion traits, and mental health. These five dimensions were derived from an independent component analysis of 109 behavioral measures (33). For each participant, propagation probability matrices were first averaged across runs and the edgewise probabilities in the resulting participant-level propagation probability matrix were used as prediction features.

Prior to prediction analysis, we controlled for potential nuisance factors, including age and head motion, in a leakage-free manner. Head motion was quantified as mean framewise displacement averaged across the four runs (34). Specifically, within each cross-validation fold, the effects of age and head motion were estimated separately on each propagation feature and the target behavior score using the only training set. The fitted general linear models were then applied to the held-out data to obtain residualized features and behavioral scores. In addition, edgewise feature scaling and univariate feature selection were performed within the training set of each fold. Features were selected based on their correlations with the residualized target score, using a threshold of $p < 0.01$. The same scaling parameters and selected feature set were then applied to the held-out participants. A linear SVR model was trained on the selected features and subsequently applied to the held-out participants for prediction. The SVR model was implemented in MATLAB using fitrsvm with a linear kernel, box constraint = 1, and epsilon = 0.1. Prediction accuracy was quantified as Pearson's correlation between predicted and observed residual scores across all out-of-fold predictions.

Statistical significance of prediction performance was assessed using permutation testing. For each behavioral dimension, the original target scores were randomly shuffled across participants 10,000 times while preserving the cross-validation fold assignments. For each permutation, the identical leakage-free pipeline was repeated, including covariate regression, feature scaling, feature selection, SVR model training, and prediction of held-out participants. This procedure generated a null distribution of prediction accuracies, from which permutation $p$ values were computed. Multiple comparisons across the five behavioral dimensions were corrected using the FDR procedure (13).

To assess the predictive contributions of different regions and systems, edge-level contribution weights were summarized at both the regional and system levels. For each cross-validation fold, edges selected as features were assigned a unit contribution weight, whereas unselected edges were assigned a weight of zero. For each edge, contribution weights were averaged across the 10 folds. Because the propagation probability matrix was directed, edge weights were summarized separately by propagation direction. For each nodal region, we separately calculated the mean weights of edges originating from and terminating in that node and then averaged these two values to quantify its overall contribution. At the system level, within- and between-system contributions were obtained by averaging edge contribution weights within each system and between each ordered pair of systems, respectively. To identify system pairs with relatively high predictive contributions, each system-level contribution weight was compared against the distribution of weights across all system pairs. System pairs with weights greater than one standard deviation above the across-pair mean were marked with asterisks in the visualization.

**Validation analysis**

To assess the robustness and reproducibility of our findings, we conducted a series of sensitivity analyses in the main (HCP-7T) dataset and cross-dataset replication in the independent MSC dataset. In the HCP-7T dataset, we varied propagation probability network construction parameters, data preprocessing strategies, and head motion control, recomputed propagation probability matrices, and repeated the main analyses. The relevant results were compared to main results (temporal window = 5 s; event threshold = 1 SD) to assess the reliability. For cross-dataset replication, we applied the same analysis framework to the MSC data to assess the reproducibility and generalizability of the main findings.

(i) Influence of event-detection and temporal-window parameters. We recomputed group-level propagation probability matrices across different combinations of ignition event thresholds (0.8 to 2.0 SD in steps of 0.2) and temporal windows (3 to 8 s in steps of 1 s). We calculated Pearson's correlations between the matrices obtained under each parameter combination and the group-level matrix from the main analysis (ignition event threshold = 1 SD and temporal window = 5 s). We further repeated the main analyses using an alternative threshold-window combination (event threshold = 2.0 SD and temporal window = 3 s).

(ii) Influence of regional hemodynamic response delays. Regional differences in hemodynamic responses may bias propagation probability estimates (35-37). To reduce this potential influence, we applied the rsHRF toolbox (37) to estimate node-specific resting-state hemodynamic response functions (HRFs) and perform node-wise deconvolution. Propagation probability matrices were then reconstructed from the deconvolved time series using the same ignition-event definition as in the main analysis, with ignition events defined as upward crossings of the node-specific z-scored signal at the predefined threshold (1 SD). We then repeated the main analyses using the resulting matrices.

(iii) Influence of head motion control. We applied a more stringent head motion control strategy by retaining participants with low head motion (mean FD across the four runs < 0.30 mm; 143 participants). We recomputed propagation probability matrices and repeated the main analyses.

(iv) Influence of brain parcellation. We repeated the main analysis pipeline with brain nodes defined using parcellations at different spatial resolutions (i.e., Schaefer-200, Schaefer-400).

(v) Cross-dataset reproducibility. To assess the reproducibility and generalizability of the main findings, we applied the same analysis pipeline to the MSC dataset. Because of the longer repetition time in MSC, we set the temporal window to 2 TRs (4.4 s; TR = 2.2 s) to approximate the 5-s window used in the main analysis.

### Supplementary Results

#### Validation Results

Across event-threshold and temporal-window combinations, the resulting group-level propagation probability matrices were highly similar to the main-analysis matrix (all *r*s > 0.941), and analyses using the alternative threshold-window combination replicaed the main findings (Fig. S4). The hemodynamic-delay-corrected matrix was also highly similar to the main-analysis matrix ($r = 0.993$), and subsequent analyses replicated the main findings (Fig. S5). Restricting the analysis to low-motion participants yielded propagation probability matrices and derived results consistent with those of the main analysis (Fig. S6). These results suggest that the main findings were not primarily attributable to specific matrix construction parameters, regional differences in hemodynamic delays, or head motion.

The main findings were also largely preserved across the Schaefer-200, Schaefer-400 parcellations (Figs. S7 and S8), supporting the robustness of the propagation probability framework across spatial resolutions. Nevertheless, subtle deviations were observed across different spatial resolutions. In the Schaefer-200 parcellation, regional propagation preferences showed modest differences in medial frontal and parietal regions (Fig. S7). These deviations may reflect the coarser spatial resolution of this parcellation, which could spatially blur heterogeneous vertex-level signals within large parcels and thereby weaken suprathreshold ignition events. As a result, regional estimates of outgoing and incoming propagation tendencies may become less spatially specific in regions with heterogeneous functional organization. Thus, although the overall propagation architecture was preserved, fine-grained regional propagation preferences should be interpreted with caution when using coarser parcellations.

Finally, the independent MSC dataset reproduced the main propagation features observed in the HCP-7T dataset, supporting cross-dataset reproducibility (Fig. S9). Specifically, the MSC analysis replicated the directed propagation organization, broad system-level architecture, and gradient-aligned canonical propagation pathways observed in the main dataset. For the canonical pattern of cluster 2, the propagation steps showed a significant negative correlation with the principal functional gradient and also exhibited deviations in the somatomotor and visual networks. These results were consistent with the main findings that the two canonical propagation patterns were not simple mirror images of each other. However, the correlations between canonical propagation pathways and the principal functional gradient were relatively weaker in MSC than in HCP-7T. This attenuation may be partly attributable to the smaller sample size of the MSC dataset and differences in acquisition parameters, including lower spatial resolution and longer TR, which may reduce the spatial and temporal precision of propagation estimates derived from spontaneous functional activity.

### Supplementary Figures

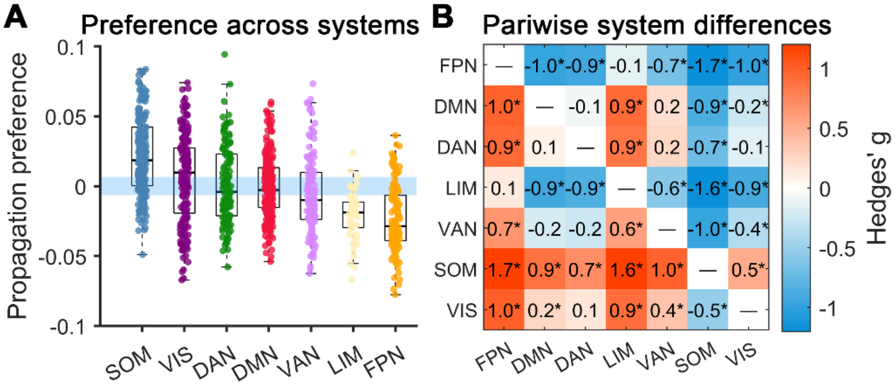


**Fig. S1.** Functional system-level differences in propagation preference. (A) Distribution of node-wise propagation preference across the seven functional systems. Propagation preference was defined for each node as the ratio of mean outgoing to mean incoming propagation probabilities and residualized by regressing out node-wise intrinsic ignition event counts. Each dot represents a nodal region, and boxplots summarize distributions across nodes within each functional system. Functional systems were ordered by their mean propagation preference. The horizontal reference band at zero indicates balanced outgoing and incoming propagation. (B) Intersystem differences in propagation preference. Each element represents Hedges' g (small-sample-corrected standardized mean difference) for pairwise comparisons of node-wise propagation preference between systems (row versus column). Positive $g$ values ($g_{ij} > 0$) indicate higher propagation preference in system $i$ than in system $j$. Asterisks indicate significant differences based on Games-Howell post hoc tests with FDR correction across all 21 comparisons ($p_{corr} < 0.05$). DAN, dorsal attention network; VIS, visual network; DMN, default-mode network; SOM, somatomotor network; FPN, frontoparietal network; VAN, ventral attention network; LIM, limbic network.

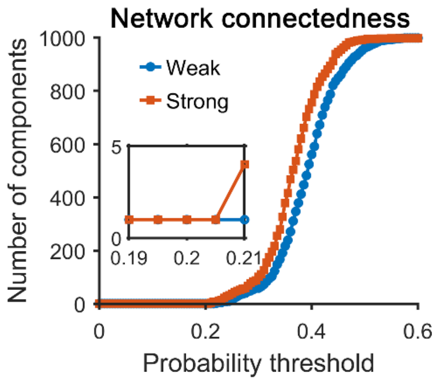


**Fig. S2.** Threshold-dependent connectedness of the propagation probability network. The number of connected components varied across probability thresholds used for network thresholding. At each threshold, edges with propagation probabilities greater than or equal to the threshold were retained to construct a directed binary network. The inset highlights the sharp transition in the number of connected components within the threshold range of 0.19-0.21, as the propagation network shifts from a largely connected configuration to a more fragmented network.

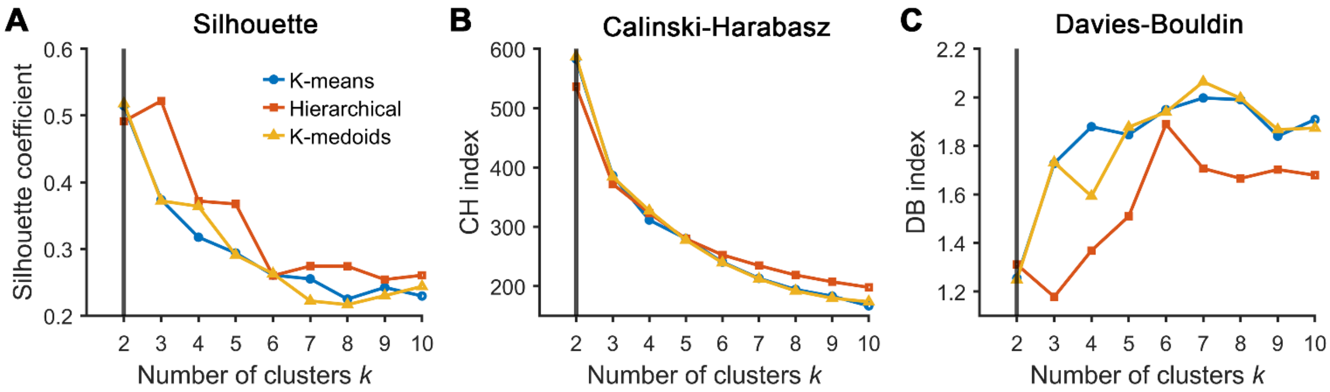


**Fig. S3.** Identification of the optimal number of clusters for canonical propagation patterns. For nodal clustering, three clustering algorithms (k-means, hierarchical clustering, and k-medoids) were separately applied to node-wise shortest-path propagation profiles. Clustering validity was evaluated as a function of the number of clusters ($k$) using three metrics: the silhouette coefficient, the Calinski-Harabasz index, and the Davies-Bouldin index. The two-cluster solution was identified in seven of the nine combinations of three clustering algorithms and three validity indices ($k = 2$; solid line), indicating a low-dimensional organization of propagation distance profiles. At the two-cluster resolution, nodal assignments were highly similar across the three algorithms (k-means vs. hierarchical: ARI = 0.72; k-means vs. k-medoids: ARI = 0.96; hierarchical vs. k-medoids: ARI = 0.73).

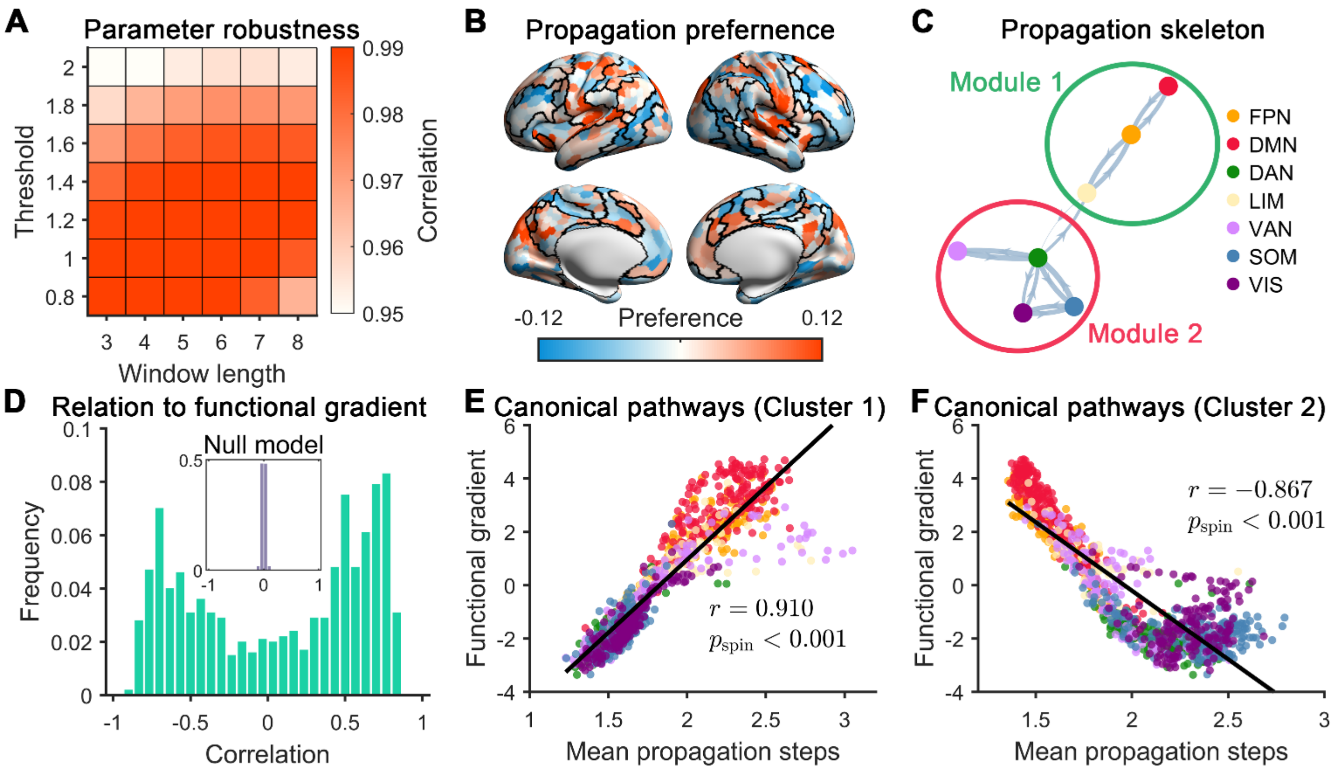


**Fig. S4.** Robustness of propagation probability structure and main findings across parameter choices. (A) Similarity of propagation probability matrices across different ignition event thresholds and temporal windows. Group-level matrices obtained across combinations of event thresholds (0.8 to 2.0 SD in steps of 0.2) and temporal windows (3 to 8 s in steps of 1 s) were compared with the matrix from the main analysis using edge-wise Pearson's correlations. Each element represents the correlation for one combination of event threshold and temporal window. All matrices showed high similarity to the main-analysis matrix (all *r*s > 0.94), indicating a robust propagation structure across parameter settings. (B-F) Replication of the main findings using an alternative parameter combination (threshold = 2.0 SD; temporal window = 3 s). (B) Spatial distribution of regional propagation preferences, corrected for regional differences in ignition event counts. (C) Modular organization of the functional system-level propagation skeleton. (D) Bimodal distribution of regional propagation-gradient associations. For each seed region, the association was quantified using Spearman's correlation between its shortest-path propagation distances to all target regions and the principal functional gradient scores of the corresponding target regions. The inset shows a null distribution obtained by permuting seed-specific shortest-path propagation distances across target regions. (E) Spatial correlation between the canonical propagation profile and the principal functional gradient for cluster 1. (F) Spatial correlation between the canonical propagation profile and the principal functional gradient for cluster 2. The statistical significance of the spatial correlations in (E) and (F) was assessed using spin-based spatial permutation tests (see *Supplementary Methods*). Together, these results demonstrate the robustness of both the propagation probability structure and the derived propagation patterns, even under the parameter combination showing the lowest similarity to the main analysis (threshold = 2.0 SD; temporal window = 3 s). DAN, dorsal attention network; VIS, visual network; DMN, default-mode network; SOM, somatomotor network; FPN, frontoparietal network; VAN, ventral attention network; LIM, limbic network.

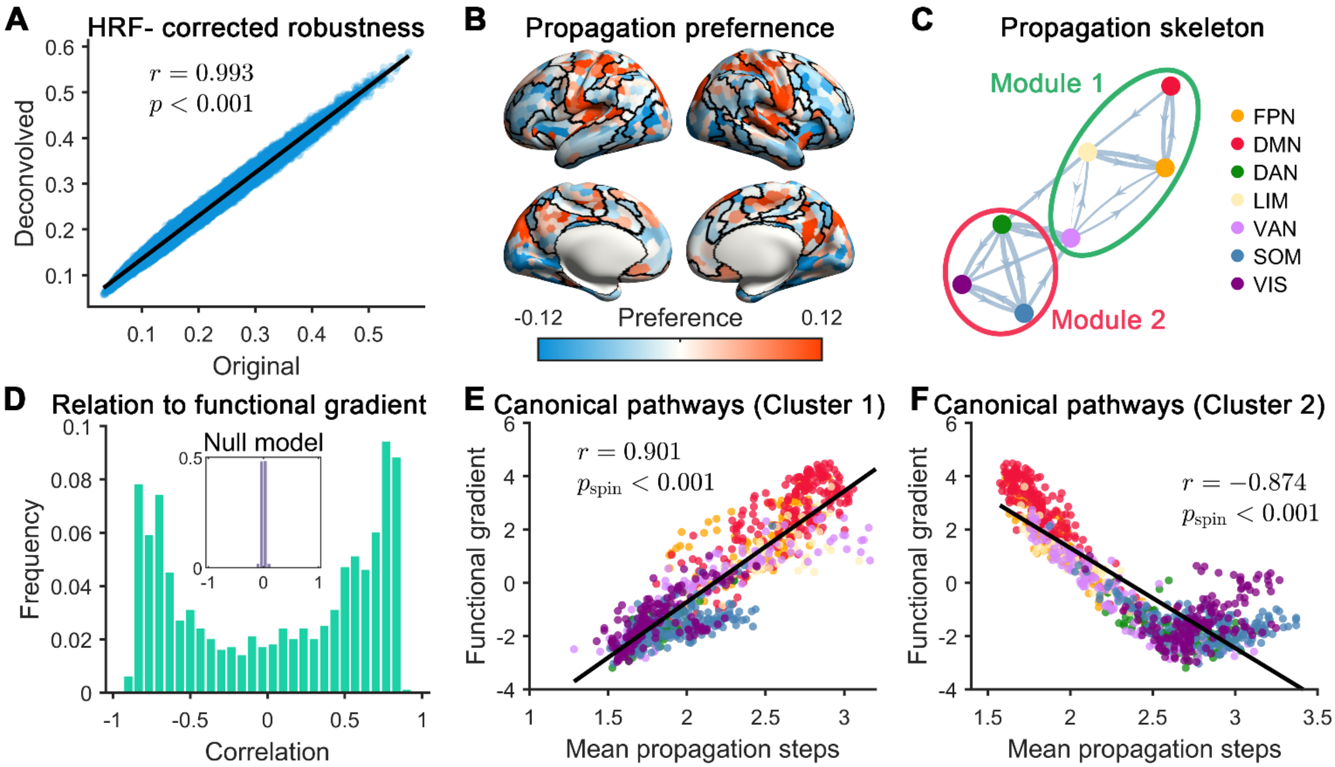


**Fig. S5.** Robustness of the ignition event propagation structure after correction for regional differences in hemodynamic delays. (A) Similarity between the group-level propagation probability matrices obtained without and with HRF correction. The high similarity between the two matrices indicates that the overall propagation structure is preserved after correcting for regional hemodynamic variability. (B-F) Replication of the main results using the group-level propagation probability matrix corrected for regional hemodynamic delays. (B) Spatial distribution of regional propagation preferences, corrected for regional differences in ignition event counts. (C) Modular organization of the functional system-level propagation skeleton. (D) Bimodal distribution of regional propagation-gradient associations. For each seed region, the association was quantified using Spearman's correlation between its shortest-path propagation distances to all target regions and the principal functional gradient scores of the corresponding target regions. The inset shows a null distribution obtained by permuting seed-specific shortest-path propagation distances across target regions. (E) Spatial correlation between the canonical propagation profile and the principal functional gradient for cluster 1. (F) Spatial correlation between the canonical propagation profile and the principal functional gradient for cluster 2. The statistical significance of the spatial correlations in (E) and (F) was assessed using spin-based spatial permutation tests (see *Supplementary Methods*). DAN, dorsal attention network; VIS, visual network; DMN, default-mode network; SOM, somatomotor network; FPN, frontoparietal network; VAN, ventral attention network; LIM, limbic network.

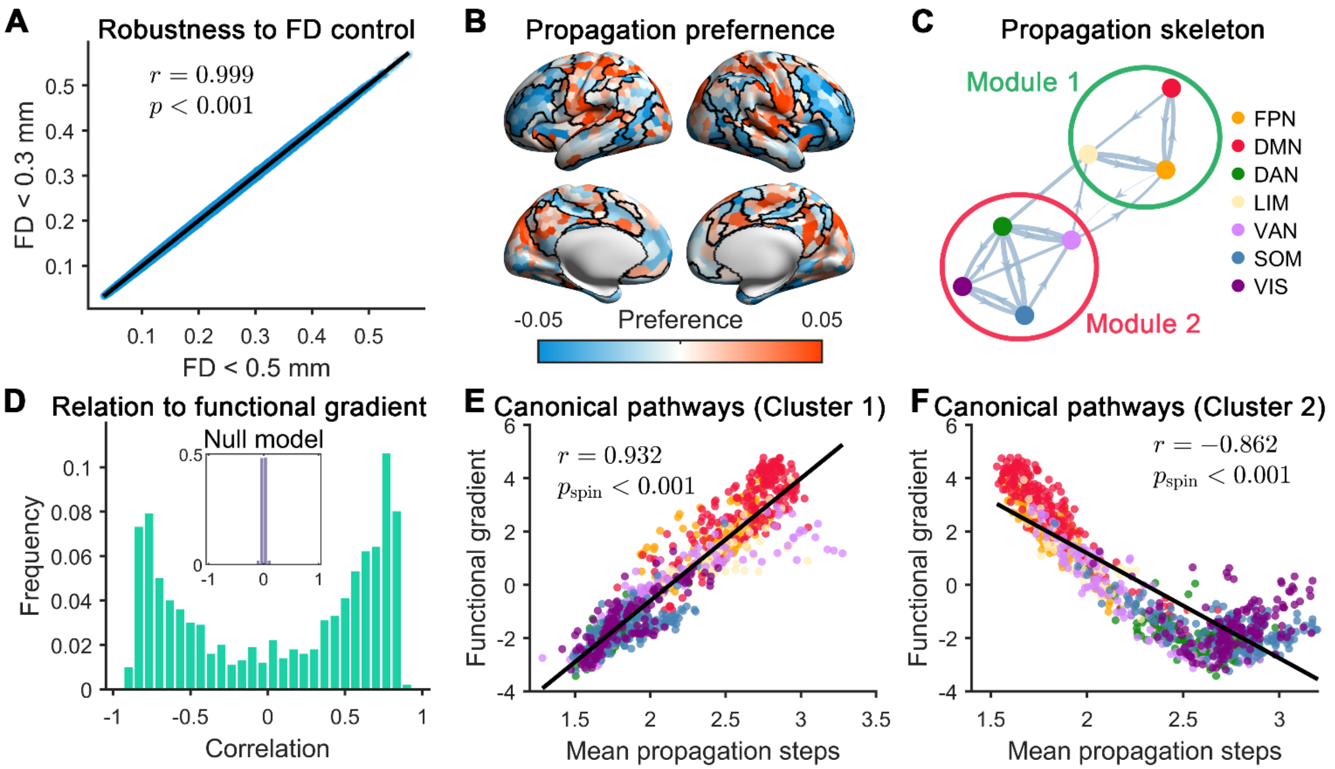


**Fig. S6.** Robustness of the ignition event propagation structure under more stringent head-motion control. (A) Similarity between the propagation probability matrix derived from a low-motion subsample (mean framewise displacement across runs < 0.30 mm) and the main-analysis matrix. High similarity indicates that the overall propagation structure is preserved after restricting the sample to low-motion participants. (B-F) Replication of the main findings in the low-motion subsample. (B) Spatial distribution of regional propagation preferences, corrected for regional differences in ignition event counts. (C) Modular organization of the functional system-level propagation skeleton. (D) Bimodal distribution of regional propagation-gradient associations. For each seed region, the association was quantified using Spearman's correlation between its shortest-path propagation distances to all target regions and the principal functional gradient scores of the corresponding target regions. The inset shows a null distribution obtained by permuting seed-specific shortest-path propagation distances across target regions. (E) Spatial correlation between the canonical propagation profile and the principal functional gradient for cluster 1. (F) Spatial correlation between the canonical propagation profile and the principal functional gradient for cluster 2. The statistical significance of the spatial correlations in (E) and (F) was assessed using spin-based spatial permutation tests (see *Supplementary Methods*). DAN, dorsal attention network; VIS, visual network; DMN, default-mode network; SOM, somatomotor network; FPN, frontoparietal network; VAN, ventral attention network; LIM, limbic network.

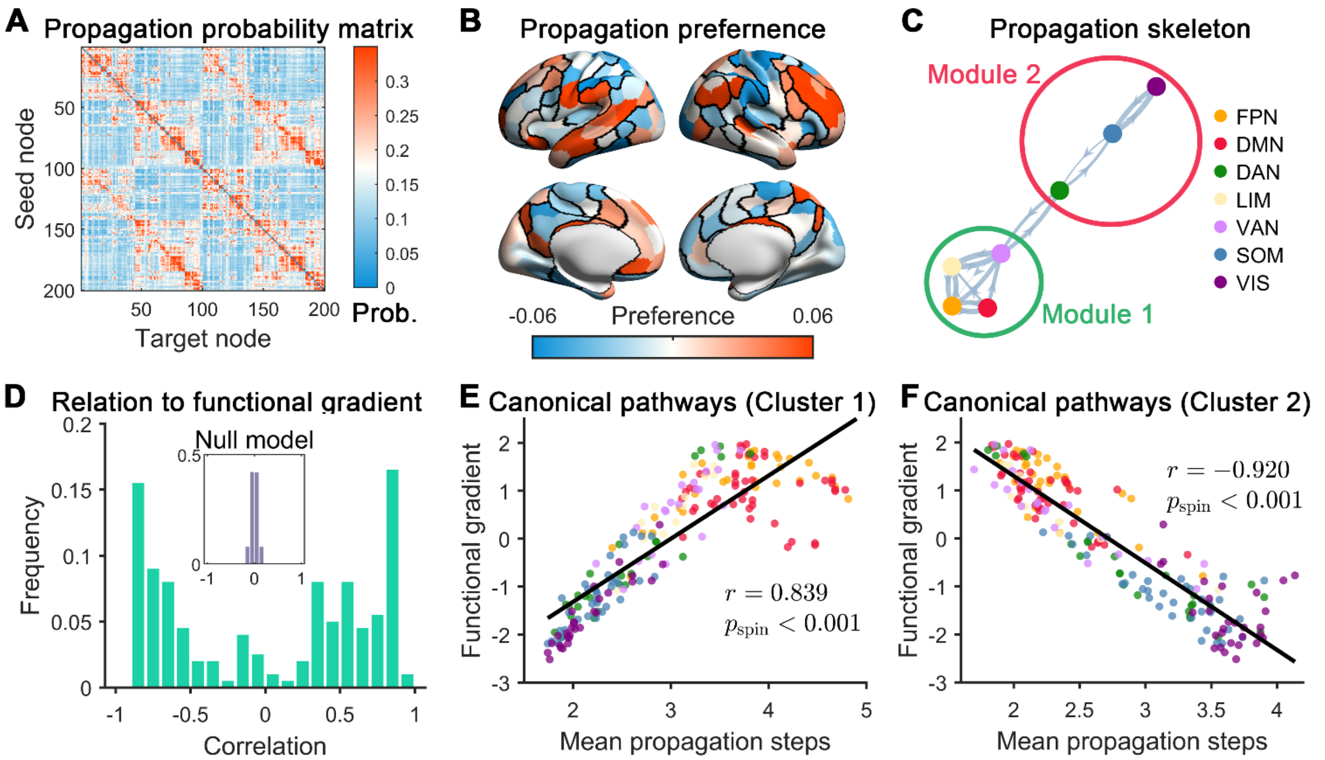


**Fig. S7.** Robustness of the propagation structure using the Schaefer 200 brain parcellation. (A) Group-level propagation probability matrix constructed using the Schaefer-200 atlas. (B-F) Replication of the main findings using the Schaefer-200 parcellation. (B) Spatial distribution of regional propagation preferences, corrected for regional differences in ignition event counts. (C) Modular organization of the functional system-level propagation skeleton. (D) Bimodal distribution of regional propagation-gradient associations. For each seed region, the association was quantified using Spearman's correlation between its shortest-path propagation distances to all target regions and the principal functional gradient scores of the corresponding target regions. The inset shows a null distribution obtained by permuting seed-specific shortest-path propagation distances across target regions. (E) Spatial correlation between the canonical propagation profile and the principal functional gradient for cluster 1. (F) Spatial correlation between the canonical propagation profile and the principal functional gradient for cluster 2. The statistical significance of the spatial correlations in (E) and (F) was assessed using spin-based spatial permutation tests (see *Supplementary Methods*). DAN, dorsal attention network; VIS, visual network; DMN, default-mode network; SOM, somatomotor network; FPN, frontoparietal network; VAN, ventral attention network; LIM, limbic network.

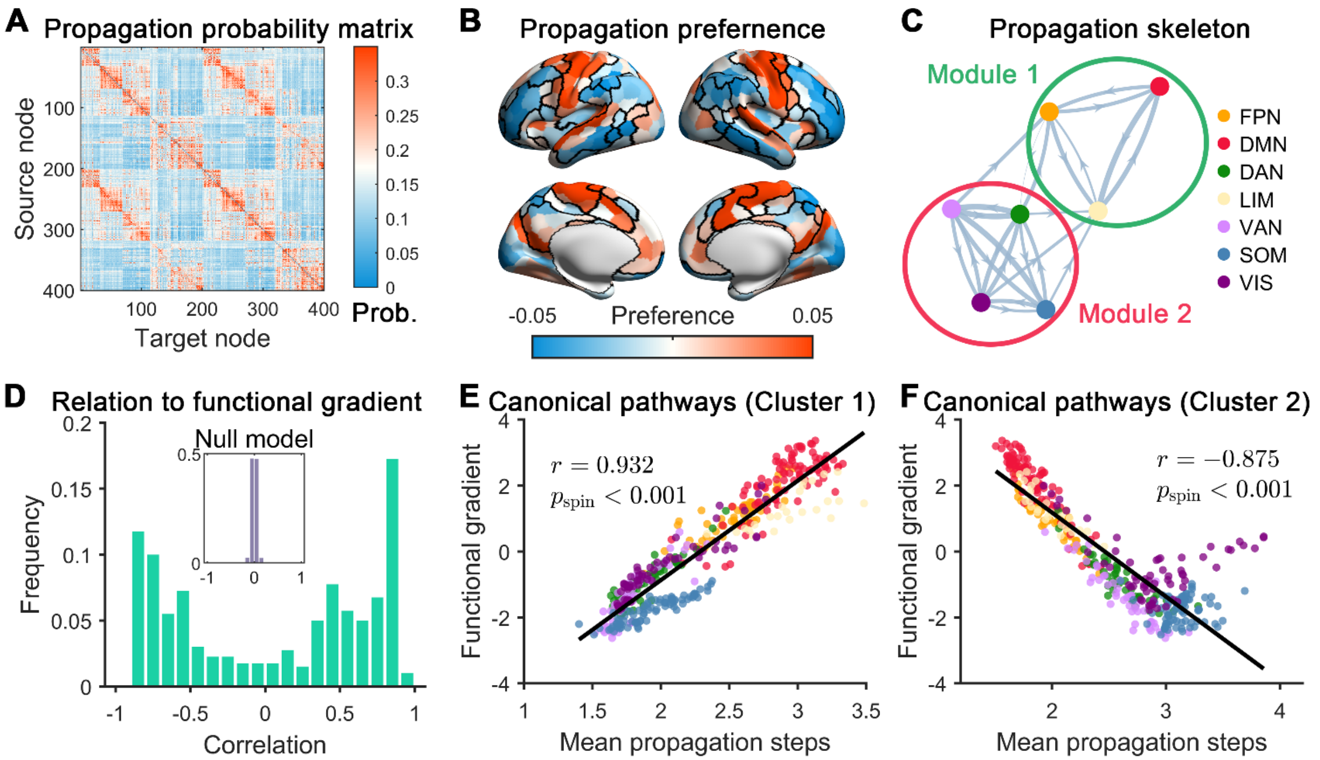


**Fig. S8.** Robustness of the propagation structure using the Schaefer-400 brain parcellation. (A) Group-level propagation probability matrix constructed using the Schaefer-400 atlas. (B-F) Replication of the main findings using the Schaefer-400 parcellation. (B) Spatial distribution of regional propagation preferences, corrected for regional differences in ignition event counts. (C) Modular organization of the functional system-level propagation skeleton. (D) Bimodal distribution of regional propagation-gradient associations. For each seed region, the association was quantified using Spearman's correlation between its shortest-path propagation distances to all target regions and the principal functional gradient scores of the corresponding target regions. The inset shows a null distribution obtained by permuting seed-specific shortest-path propagation distances across target regions. (E) Spatial correlation between the canonical propagation profile and the principal functional gradient for cluster 1. (F) Spatial correlation between the canonical propagation profile and the principal functional gradient for cluster 2. The statistical significance of the spatial correlations in (E) and (F) was assessed using spin-based spatial permutation tests (see *Supplementary Methods*). DAN, dorsal attention network; VIS, visual network; DMN, default-mode network; SOM, somatomotor network; FPN, frontoparietal network; VAN, ventral attention network; LIM, limbic network.

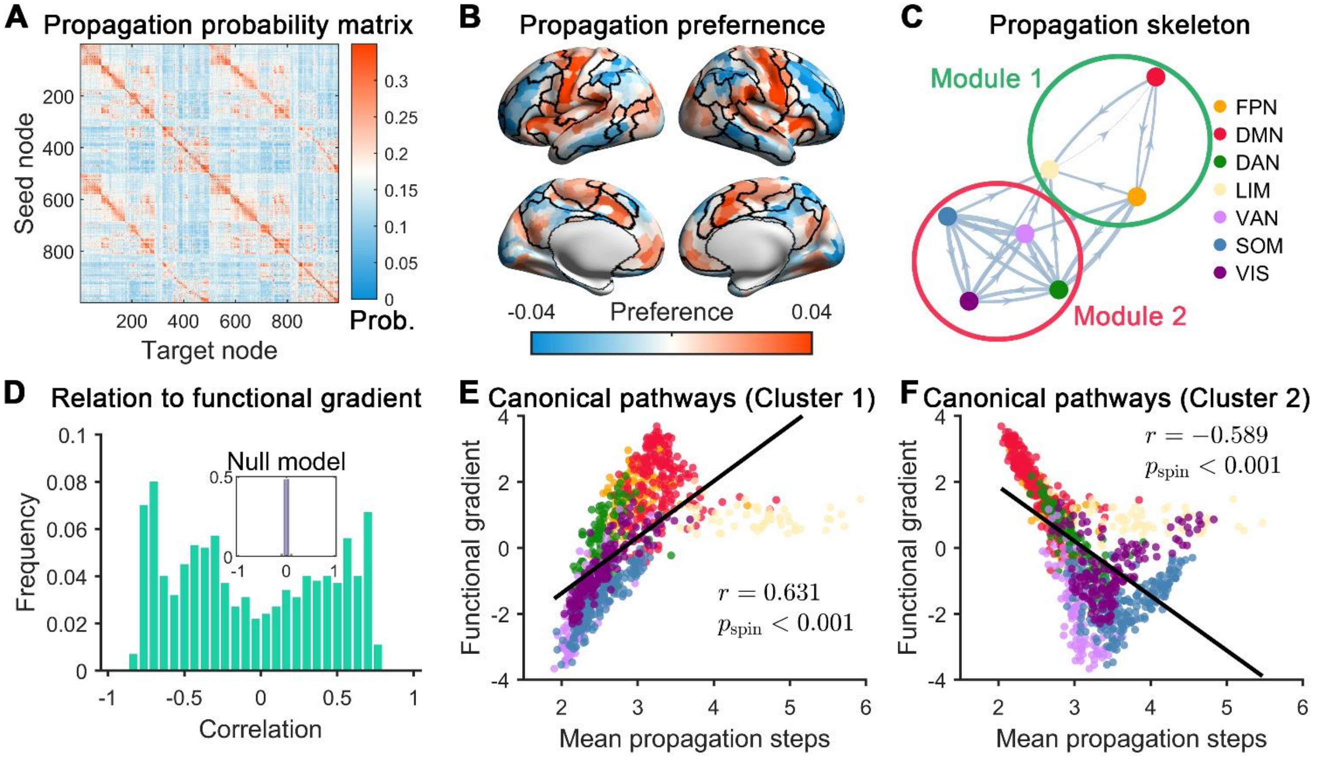


**Fig. S9.** Cross-dataset reproducibility of the propagation structure in an independent Midnight Scan Club dataset. (A) Group-level propagation probability matrix constructed using the Midnight Scan Club dataset. The matrix was defined using parameters (ignition threshold = 1SD and temporal window = 4.4 s) that approximate those used in the main analysis. (B-F) Replication of the main findings in the independent dataset. (B) Spatial distribution of regional propagation preferences, corrected for regional differences in ignition event counts. (C) Modular organization of the functional system-level propagation skeleton. (D) Bimodal distribution of regional propagation-gradient associations. For each seed region, the association was quantified using Spearman's correlation between its shortest-path propagation distances to all target regions and the principal functional gradient scores of the corresponding target regions. The inset shows a null distribution obtained by permuting seed-specific shortest-path propagation distances across target regions. (E) Spatial correlation between the canonical propagation profile and the principal functional gradient for cluster 1. (F) Spatial correlation between the canonical propagation profile and the principal functional gradient for cluster 2. The statistical significance of the spatial correlations in (E) and (F) was assessed using spin-based spatial permutation tests (see *Supplementary Methods*). DAN, dorsal attention network; VIS, visual network; DMN, default-mode network; SOM, somatomotor network; FPN, frontoparietal network; VAN, ventral attention network; LIM, limbic network.
